# Elevated hydrostatic pressure modulates endothelial junctional mechanotransduction through VE-cadherin remodelling and altered association with YAP1, EPS8: an endothelium-on-chip study

**DOI:** 10.64898/2026.08.31.748221

**Authors:** P. Vasanthi Bathrinarayanan, Thomas Abadie, D. Vigolo, M. J. H. Simmons, L. M. Grover

## Abstract

Endothelial dysfunction is a hallmark of numerous vascular pathologies and is strongly influenced by mechanobiological forces within the vascular microenvironment. While the effects of shear stress have been extensively investigated, the mechanisms by which elevated hydrostatic pressure regulates endothelial junctional organisation remain sparsely investigated. Here, we employed a microfluidic platform to investigate the combined effects of low shear stress (1.4 dyne/cm^2^) and elevated hydrostatic pressure (∼3972 Pa) on endothelial junctional dynamics. Elevated hydrostatic pressure induced marked remodelling of VE-cadherin junctions, characterised by formation of serrated, finger-like structures accompanied by increased YAP1 nuclear localisation and reduced YAP1-VE-cadherin cytoplasmic colocalisation compared to shear stress alone conditions. Further, elevated hydrostatic pressure also demonstrated an increase in cytoplasmic accumulation of EPS8, an actin adaptor protein, and increased cytoplasmic EPS8-VE-cadherin colocalisation. These observations were accompanied by functional changes marked by increased endothelial permeability, and enhanced THP-1 monocyte adhesion, thus suggesting activation of mechanosensitive pathways linked to dynamic junctional reorganisation. Inhibition of PI3K at elevated hydrostatic pressure exhibited a thin VE-cadherin patterning and increased cytoplasmic EPS8-VE-cadherin colocalisation, thus demonstrating a prominent role for PI3K signalling in regulating the junction organisation. Interestingly, Piezo-1 activation using Yoda1 produced context-dependent effects. Under shear stress alone, Yoda1 promoted YAP1 nuclear translocation, reduced YAP1-VE-cadherin colocalisation, increased endothelial permeability but strikingly did not impact THP-1 adhesion compared to shear stress alone conditions. In contrast, under elevated hydrostatic pressure conditions, Yoda1 significantly reduced both endothelial permeability and THP-1 adhesion while increasing YAP1-VE-cadherin colocalisation and decreasing YAP1 nuclear accumulation. Collectively, these findings identify a previously underappreciated elevated hydrostatic pressure-Piezo-1-PI3K signalling axis that regulates endothelial barrier integrity and pro-adhesive endothelial activation through coordinated regulation of VE-cadherin, YAP1, and EPS8. These results highlight elevated hydrostatic pressure as a unique mechanobiological stimulus, distinct from that of shear stress alone and provide novel insights into mechanisms underlying microvascular dysfunction.

## 2 Introduction

Endothelial mechanotransduction plays a critical role in vascular homeostasis and in the pathogenesis of diseases [1] including atherosclerosis [2], glaucoma [3], pulmonary oedema [4], and acute compartment syndrome [5]. Endothelial cells continuously sense and respond to mechanical cues within the vascular microenvironment, translating physical forces into biochemical signalling programmes that regulate proliferation, migration, cytoskeletal organisation, permeability and inflammation [1, 6]. Endothelial dysfunction involves dysregulation of these mechanosensitive processes and can contribute to tissue oedema, impaired permeability, leukocyte recruitment, ultimately contributing to the pathogenesis of various tissues [7, 8]. Endothelial cells experience three major forces, namely fluid shear stress generated by blood flow, cyclic stretch arising from pulsatile vessel wall deformation, and hydrostatic pressure imposed by intraluminal pressure [9]. Although these forces act upon endothelial cells simultaneously in vivo, they are often studied in isolation which limits the understanding of how these combined mechanical cues regulate endothelial phenotype.

Among the different mechanical stimuli, the influence of shear stress on endothelial functions has been extensively investigated [10]. Past studies have established that laminar shear stress promotes an atheroprotective endothelial phenotype [11] by inducing actin alignment in the direction of flow [12], enhancing nitric oxide production [13], maintaining barrier function [14] and suppressing inflammatory activation [15]. In contrast, disturbed and oscillatory shear stress have been shown to promote endothelial dysfunction through increased reactive oxygen species production, actomyosin stress fibre formation, junctional remodelling, and enhanced expression of adhesion molecules that support leukocyte recruitment [16, 17]. While the majority of the studies have focused on the atheroprotective shear stress range of ∼10-25 dyne/cm^2^ to represent the quiescent endothelial phenotype of a straight arterial vessel [18, 19], there is a requirement to understand the impact of very low shear stresses (≤5 dyne/cm^2^) on endothelial cells [20, 21] especially considering their impact on junctional regulation [22–24] and subsequent atherogenic pathology development.

In contrast to shear stress and cyclic stretch, the effects of hydrostatic pressure on endothelial junctional dynamics have been sparsely investigated. Brockhaus et al (2020) reported that elevated hydrostatic pressure (30-60 mmHg) downregulated adherens junction VE-cadherin, vascular endothelial growth factor 1 (VEGF1), and intermediate filament vimentin whereas the tight junction protein ZO-1 was upregulated in retinal endothelial cells [25]. Similarly, Ohashi et al (2007) showed that junctional VE-cadherin was downregulated at elevated hydrostatic pressure (50-150 mmHg) resulting in increased proliferation due to loss of contact inhibition [26]. Other emerging studies have also shown the potent mechanotransductive effect of elevated hydrostatic pressure on angiogenesis [27], endothelial tube formation [28], extracellular matrix proteins fibronectin and laminin expression levels [29] and cytoskeletal reorganisation [30]. Despite providing valuable insights, many of these studies were performed under static culture conditions, where endothelial cells were exposed to elevated pressure in the absence of shear stress. To increase the physiological relevance and translatability of studies, there is a requirement to examine the combined influence of shear stress and hydrostatic pressure on endothelial junctional organisation, barrier function, and inflammatory activation.

Several mechanosensory proteins have been implicated in vascular mechanotransduction including G-protein coupled receptors, glycocalyx, ion channels and primary cilia [31, 32]. Piezo-1 is a cation-permeable mechanosensitive channel expressed in endothelial cells and plays a key role in flow mediated force transmission [33, 34]. Upon activation, Piezo-1 opening leads to Ca²⁺ influx and regulation of downstream processes including cytoskeletal remodelling [35], vascular tone [36], endothelial inflammation [37, 38], and barrier function [39]. At a molecular level, Piezo-1 integrates with broader mechanotransduction networks, including phosphoinositide-3-kinase-protein kinase B/Akt (PI3K/Akt) [40, 41] and the Hippo–YAP/TAZ signalling pathways which together govern endothelial junctional organisation, barrier stability, and inflammatory responses [42, 43].

The Hippo pathway effector molecule Yes-associated protein 1 (YAP1) has been shown to act as a major transcriptional mechanotransducer whose nuclear localisation is regulated by cytoskeletal tension and junctional integrity [44, 45]. Emerging studies have shown that VE-cadherin-containing junctional complexes can restrain YAP1 activity [27, 46], whereas junctional remodelling and increased actomyosin tension can promote YAP1 nuclear accumulation [42, 47]. In addition, Epidermal growth factor receptor pathway substrate 8 (EPS8), an actin-binding adaptor protein, has emerged as a key regulator of actin polymerisation, thereby impacting cell processes such as morphogenesis [48], migration [49] and endocytosis [50]. Recently, Giampietro et al (2015) reported on a mechanobiological framework that demonstrated the influence of EPS8 and YAP1 in regulating VE-cadherin turnover and endothelial permeability [51, 52]. However, the impact of elevated hydrostatic pressure in regulating VE-cadherin, YAP1, and EPS8 junctional dynamics is still less understood.

Microfluidic organ-on-chip systems provide a powerful approach to investigate endothelial mechanobiology under controlled and physiologically relevant mechanical conditions [53]. Previous studies have demonstrated the potential of microfluidic platforms to investigate the influence of physiological and pathological flows on endothelial barrier function, calcium signalling, inflammatory activation, and mechanobiological signalling [54–56]. While these advanced platforms have been extensively used model diverse microvascular pathologies [57], precise understanding of localised junctional mechanics is still evolving. Importantly, microfluidic models permit the examination of combined mechanical forces such as shear stress, hydrostatic pressure and cyclic stretch [58, 59], thereby enabling mechanistic interrogation of force-sensitive signalling pathways at the endothelial junctions.

In this study, we employed a simple, easy-to-implement microfluidic endothelium-on-chip model to investigate the combined influence of low shear stress (1.4 dyne/cm^2^) and elevated hydrostatic pressure (∼3972 Pa) and on VE-cadherin junctional organisation, YAP1 localisation, EPS8 dynamics, junctional permeability, and THP-1 monocyte adhesion. We further examined the influence of pharmacological activation of Piezo-1 using Yoda1 and inhibition of PI3K using Wortmannin on the observed elevated pressure-dependent endothelial responses. We hypothesised that elevated hydrostatic pressure promotes endothelial junctional remodelling and pro-adhesive inflammatory phenotype through coordinated changes in VE-cadherin, YAP1, and EPS8, which in turn are influenced by Piezo-1 and PI3K signalling. In doing so, this study proposes elevated hydrostatic pressure as a distinct mechanobiological stimulus that warrants integration into the broader endothelial mechanobiology framework.

## 3 Methods

### 3.1 Cell culture

HUVECs were selected for the current study due to their ability to more faithfully recapitulate human endothelial behaviour in vivo compared to immortalized cell lines [60]. Given their high physiological relevance, HUVECs have been used as a standard, versatile in vitro platform for investigating vascular diseases and for toxicology and pharmacology studies [61]. For all experiments, HUVECs (PromoCell, Heidelberg, Germany) were used between passages 3 and 8. Cells were maintained in M199 growth medium supplemented with 10% heat-inactivated foetal bovine serum (FBS; Gibco™, Sigma-Aldrich, UK) and 3% (v/v) endothelial cell growth supplement (Cell Applications Inc., Merck, UK). The human acute monocytic leukaemia-derived THP-1 cell line (Sigma Aldrich, Merck Ltd, Dorset, UK) was maintained in complete RPMI 1640 medium (Gibco™, Sigma-Aldrich, UK) supplemented with 10% FBS, 2mM Glutamine (Sigma Aldrich, Merck Ltd, Dorset, UK) and 1% penicillin-streptomycin (Lonza, Basel, Switzerland).

### 3.2 Microfluidic device fabrication

Microfluidic devices were fabricated using standard soft-lithography processes [62]. Briefly, the channel geometry [1000 µm (w) x 100 µm (h) x 17 mm (l)] was designed in AutoCAD (Autodesk, San Francisco, US), imprinted onto a photomask (Micro Lithography Services Ltd, Chelmsford, UK) and printed onto a silicon master mould using SU-8 2075 photoresist (A-Gas Electronic Materials Limited, Rugy, UK) via standard photolithography procedures [62]. PDMS moulds from soft-lithography were produced using standard SYLGARD 184® Silicon Elastomer kit (Dow Corning, USA) by mixing the monomer and curing agent at a ratio 10:1 and casting the mixture onto the silicone mould. The mixture was then degassed using a vacuum pump to remove air bubbles and cured in an oven at 70°C for 2 h. Following curing, the PDMS devices were cut from the moulds using a scalpel, and inlet/outlet ports were punched using a 1.5mm biopsy punch (Miltex® Biopsy Punch, Medisave, Dorset, UK). The devices were then bonded to glass substrates via corona discharge treatment (Piezobrush® PZ3, Intertronics, Oxfordshire, UK) and annealed on a hot plate at 95°C for 20 min to ensure a stable, hermetic seal. For experiments, the PDMS moulds were autoclaved, treated with 70% ethanol and washed with Dulbecco’s phosphate buffered saline (DPBS) twice to ensure sterilisation.

### 3.3 Microfluidic perfusion experiments

Prior to cell seeding, the microfluidic channels were coated with 0.2% gelatine (Sigma Aldrich, Merck Ltd, Dorset, UK) and incubated at 37°C and 5% CO_2_ for 2h. The channels were washed with warm growth medium to remove any excess gelatine after which 10 µL HUVECs from a cell suspension of 10^7^ cells/mL were gradually introduced into the channels (10^5^ cells/channel) taking care that no air bubbles were formed during this process. The cell-laden microfluidic devices were incubated at 37°C and 5% CO_2_ for 2 h to ensure firm adhesion of the endothelial cells to the channel bottom. Cells were then examined for uniform monolayer formation under a brightfield microscope (EVOS M3000, Invitrogen, Fisher Scientific, Loughborough, UK) after which they were connected to the inlet/outlet tubing (Saint-Gobain Tygon Microbore Tubing, Fisher Scientific, Loughborough, UK). The inlet tubing was connected using blunt-end Luer Lock Syringe Needles (Darwin Microfluidics, Paris, France) to a 5mL Luer Lock syringe (BD Emerald, Medisave, Dorset, UK) containing warm growth media and perfusion was implemented using a syringe pump (World Precision Instruments, Hitchin, UK).

### 3.4 Shear stress and hydrostatic pressure distributions

The shear stress (SS) and hydrostatic pressure (HP) distributions across and along the channels were reported in our previous study which used a microfluidic channel of similar dimensions [63]. The cells were exposed to a flow rate of 13 µL/min and the wall shear stress distribution along the bottom wall of the channel was derived using analytical solutions (Supplementary section S1 for wall shear stress analytical equations for a rectangular channel). The mentioned flow rate produced a peak shear stress of 1.4 dyne/cm^2^ (0.14 Pa) in the central region of the channel (100 µm on either side of the centre of the channel) where the cells were imaged. Elevated hydrostatic pressure conditions were achieved by raising the outlet tube to 40 cm above the inlet. At this height, the pressure levels at the channel inlet and outlet were analytically derived [64, 65] to be 3971.18 Pa and 3924 Pa respectively (Supplementary section S1 for analytical equations governing pressure calculations and for the full range of pressure values along the length of the channel).

**Table 1.** Flow rates, shear stress and hydrostatic pressure values for the two flow conditions used in this study.

| Condition | Flow rate | Shear stress ($\text{dyne}/\text{cm}^2$) | Inlet HP (Pa) | Outlet HP (Pa) |
| --- | --- | --- | --- | --- |
| SSlow only | 13$\mu\text{L}/\text{min}$ | 1.4 | 47.1 | 0 |
| SSlow+HP | 13$\mu\text{L}/\text{min}$ | 1.4 | 3971.18 | 3924 |

### 3.5 Avidin-FITC endothelial permeability assay

The endothelial barrier function was investigated using an assay adapted from Dubrovskyi et al (2013) [66]. The assay quantifies the permeability of FITC-conjugated avidin across the endothelial monolayer to bind to biotinylated gelatine in response to different flow regimes and mechanosensitive agonists such as Yoda1. Briefly, EZ-Link NHS-LC-LC-Biotin (Fisher Scientific, Loughborough, UK) was dissolved in 0.2% gelatine solution to give a working concentration of 0.25 mg/mL. Biotin conjugation to gelatine was produced by constantly stirring the solution at room temperature for 24 h. HUVECs were cultured on biotinylated gelatine coating within microfluidic devices as described in Section 3.3 and were treated to the different flow conditions in the presence or absence of 2 µM Yoda1 for 1 h. After 1 h, the cells were washed once with DPBS after which Avidin-FITC solution (Fisher Scientific, Loughborough, UK) at a concentration of 2.5 µg/mL was introduced into the microchannel and incubated for 3 min at room temperature. Unbound Avidin-FITC was flushed out of the microchannel by washing twice with DPBS. The cells were then fixed by flowing 10% formalin neutral buffered solution (Sigma Aldrich, Merck Ltd, Dorset, UK) for 10 min at room temperature into the microchannel, washed twice with DPBS and then stained for F-actin using Phalloidin CF^®^594 (Biotium, Inc., CA, USA) for 5 mins at room temperature. Finally, the biotin bound Avidin-FITC was quantified by imaging under Olympus FLUOVIEW Spectral FV3000 confocal microscope (Olympus, Tokyo, Japan) predominantly using the 20x (C-Apochromat, 1.2 W Korr FCS M27) objectives. For image analysis, five standardized 200 × 200 µm regions of interest (ROIs) were applied to each image (3 images/condition) using the ROI Manager tool, with four ROIs positioned at the image corners and one centrally located in order to capture the mean fluorescence intensity (F.I.) of the image. Identical ROI size and positions were maintained across all images, and the F.I. was measured for each ROI. The F.I. values for each condition were then normalised to the mean control F.I. prior to statistical analysis.

### 3.6 THP-1 adhesion

HUVECs were cultured within microfluidic channels as described in Section 3.3 and subjected to specified flow conditions in the presence or absence of 2 µM Yoda1 for 45 min. For the positive control (SSlow + TNF-α), HUVECs were pre-stimulated with 10 ng/mL TNF-α for 24 h under SSlow only conditions prior to monocyte perfusion. Following treatment, the devices were transferred to a sterile stage-top incubator (Olympus FLUOVIEW Spectral FV3000, Tokyo, Japan) for live-cell time-lapse imaging. THP-1 monocytes were incubated with 2 µM CellTracker™ Deep Red dye (Invitrogen, Fisher Scientific, Loughborough, UK) in serum-free medium for 20 min at 37°C after which they were centrifuged and washed twice with growth medium to remove any unbound dye. The labelled THP-1 monocytes were then resuspended at a density of 200,000 cells/mL and perfused over the HUVEC monolayer under the specified flow conditions for 15 min. To quantify steady-state monocyte adhesion, time-lapse sequences were captured at a frame interval of 2 s (0.5 Hz) during the final 3 min of perfusion. For analysis, firmly adherent THP-1 cells were quantified from the final frame of each experimental replicate (n=3 images per condition). Adhered THP-1 were averaged per condition and normalised to control values prior to statistical analysis.

### 3.7 Immunostaining

Immunofluorescence staining was performed by flowing reagents, primary and secondary antibodies over the cells fixed within the microfluidic devices. For immunostaining, the cells were fixed by flowing 10% formalin neutral buffered solution (Sigma Aldrich, Merck Ltd, Dorset, UK) for 10 min at room temperature into the microchannel followed by incubation with 0.1% Triton-X in DPBS (Gibco, Merck Ltd, Dorset, UK) for 5 min at room temperature for permeabilization of the cell membrane. The channels were then washed 3 times using DPBS and were subsequently blocked using 1% BSA (Sigma Aldrich, Merck Ltd, Dorset, UK) diluted in DPBS containing 0.1% Tween-20 (Sigma Aldrich, Merck Ltd, Dorset, UK) for 30 min. The channels were washed thrice with DPBS and were incubated with 5 μg/ml (diluted in blocking buffer) anti-human mouse monoclonal VE-cadherin (CD144) (Invitrogen, Fisher Scientific, Loughborough, UK), anti-human rabbit polyclonal EPS8 (Novus Biologics, Bio-Techne Limited, Abingdon, UK), anti-human CoraLite® Plus 488-conjugated YAP1 rabbit polyclonal antibody (Proteintech Europe, Manchester, UK) at 4 °C overnight. The following day, the unbound primary antibodies were removed by washing the channels three times with DPBS and the cells were subsequently incubated for 1 h at room temperature with goat anti-mouse secondary antibodies conjugated with Alexa Fluor™ 488 (Invitrogen, Fisher Scientific, Loughborough, UK), goat anti-rabbit conjugated with Alexa Fluor™ 647, (Invitrogen, Fisher Scientific, Loughborough, UK) or goat anti-mouse secondary antibodies conjugated with Alexa Fluor™ 647 at a dilution of 1:1000 in blocking buffer. After 1 h, the unbound secondary antibodies were removed by washing 3 times with DPBS. The cells were then stained for nuclei using DAPI (4′,6-diamidino-2-phenylindole, Sigma Aldrich, Merck Ltd, Dorset, UK) for 5 min at room temperature. The cells were then imaged (while still within the microfluidic device) using a Carl Zeiss LSM880 scanning confocal microscope (Carl Zeiss AG, Germany) predominantly using the 40× (C-Apochromat, 1.2 W Korr FCS M27) and 63× (C-Apochromat, 1.2 W Korr M27) objectives.

### 3.8 Image analysis

All the raw images were processed (brightness/contrast adjustments) either using FIJI (US National Institutes of Health) or Zeiss Zen software (Carl Zeiss AG, Germany). For cell area, cell circularity and aspect ratio, cell outlines were drawn using the freehand selection tool and analysed using FIJI. For all other colocalisation analysis, customized pipelines on CellProfiler 4.2.6 (Broad Institute, Massachusetts Institute of Technology, USA) were used to automate image analysis. Briefly, nuclei and EPS8 or YAP1 raw images, designated as primary and secondary objects, respectively, were thresholded using a minimum cross-entropy method. Cytoplasm fluorescence intensity was derived by subtracting the primary object (DAPI stain) from the secondary object (EPS8 or YAP1 stain). EPS8 cytoplasmic area, EPS8-VE-cadherin, YAP1-VE-cadherin, YAP1-DAPI colocalisation were quantified using the “Measure ObjectIntensity” modules. The pipelines used for image analysis on CellProfiler can be provided upon reasonable request.

### 3.9 Agonists and inhibitors

For Piezo-1 cation-permeable mechanosensitive channels activation experiments, cells were treated to growth media containing 2 µM Yoda1 (Tocris bioscience, Bio-Techne, UK). For PI3K inhibition experiments, cells were treated with growth media containing 1.25 µM Wortmannin (Tocris bioscience, Bio-Techne, UK). After exposure, cells were washed thrice with DPBS and were fixed for immunostaining.

### 3.10 Statistical analysis

All statistical analysis was performed using GraphPad Prism10.3.1 (GraphPad, Boston, US). One-way ANOVA along with Tukey’s post-hoc test was used to compare between different conditions. Statistical significance was set as follows: *p < 0.05, **p < 0.01, ***p < 0.001, and ****p < 0.0001. Statistical tests and relative p values are indicated in each figure legend. Unless stated elsewhere, all experiments were performed with at least three independent biological replicates (three microfluidic channels).

## 4 Results

### 4.1 Elevated hydrostatic pressure leads to increased YAP1 nuclear translocation and reduced association with VE-cadherin

YAP1 closely associates with junctional VE-cadherin during resting state whereas disruption of VE-cadherin in response to mechanical stress results in YAP1 nuclear translocation and subsequent transcription of pro-inflammatory genes [44, 46]. In the current study, HUVECs were exposed to the different flow conditions for 1 h in the presence or absence of Piezo-1 agonist Yoda1 (2 µM). Using immunostaining procedure described in methods section 3.7, VE-cadherin expression pattern, YAP1 nuclear localisation and YAP1-VE-cadherin colocalisation were analysed.

When exposed to the different flow conditions, SSlow only and CTRL conditions demonstrated a continuous, linear VE-cadherin expression pattern at cell-cell junctions as shown in Figure 1 (B) and (R) respectively (linear VE-cadherin pattern denoted by white arrows) whereas SSlow+Yoda1 demonstrated a linear but intermittently disrupted VE-cadherin pattern at the cell-cell junctions as shown in Figure 1 (F) (disrupted VE-cadherin junctions denoted by yellow arrows).

**Figure 1.**
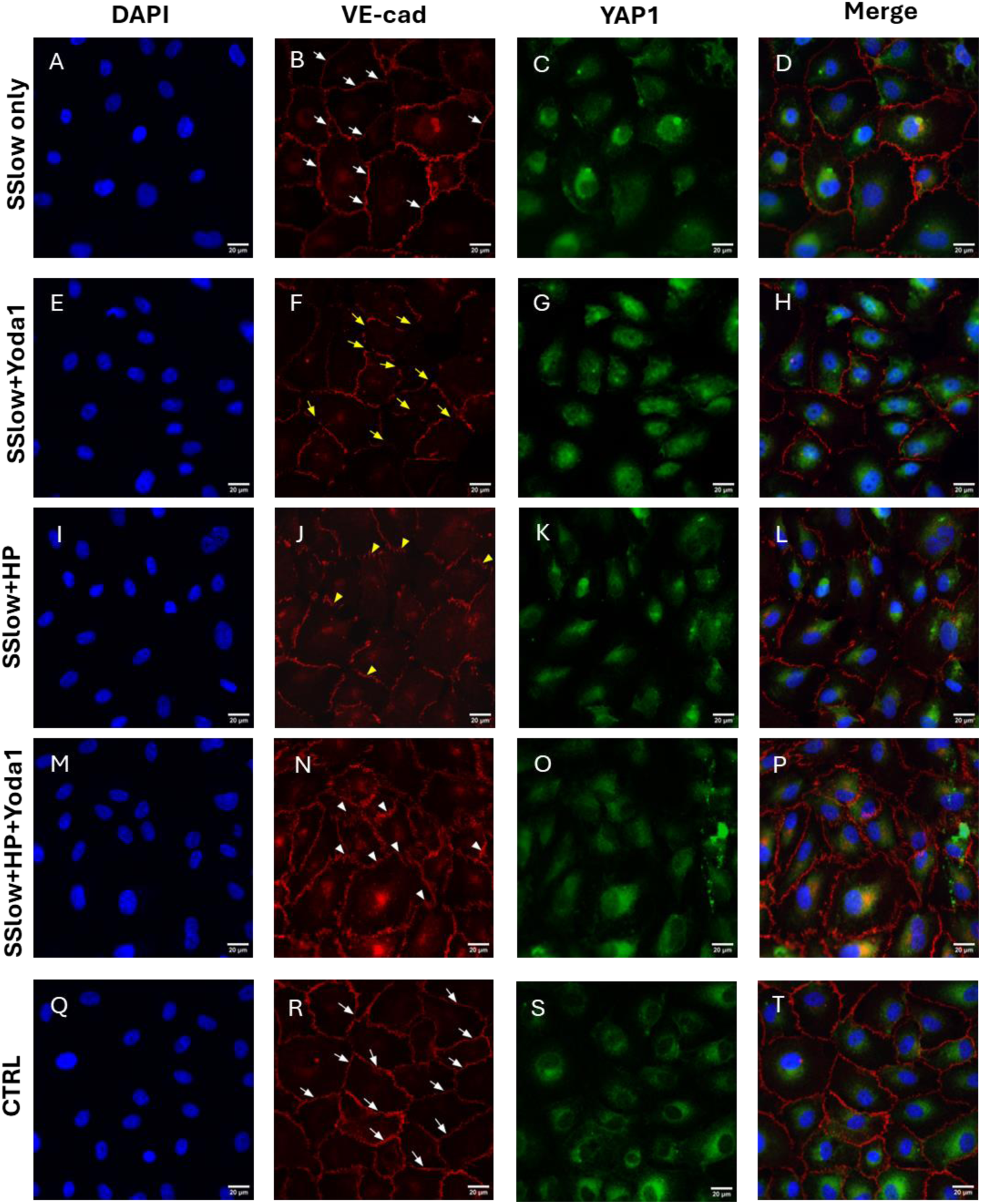
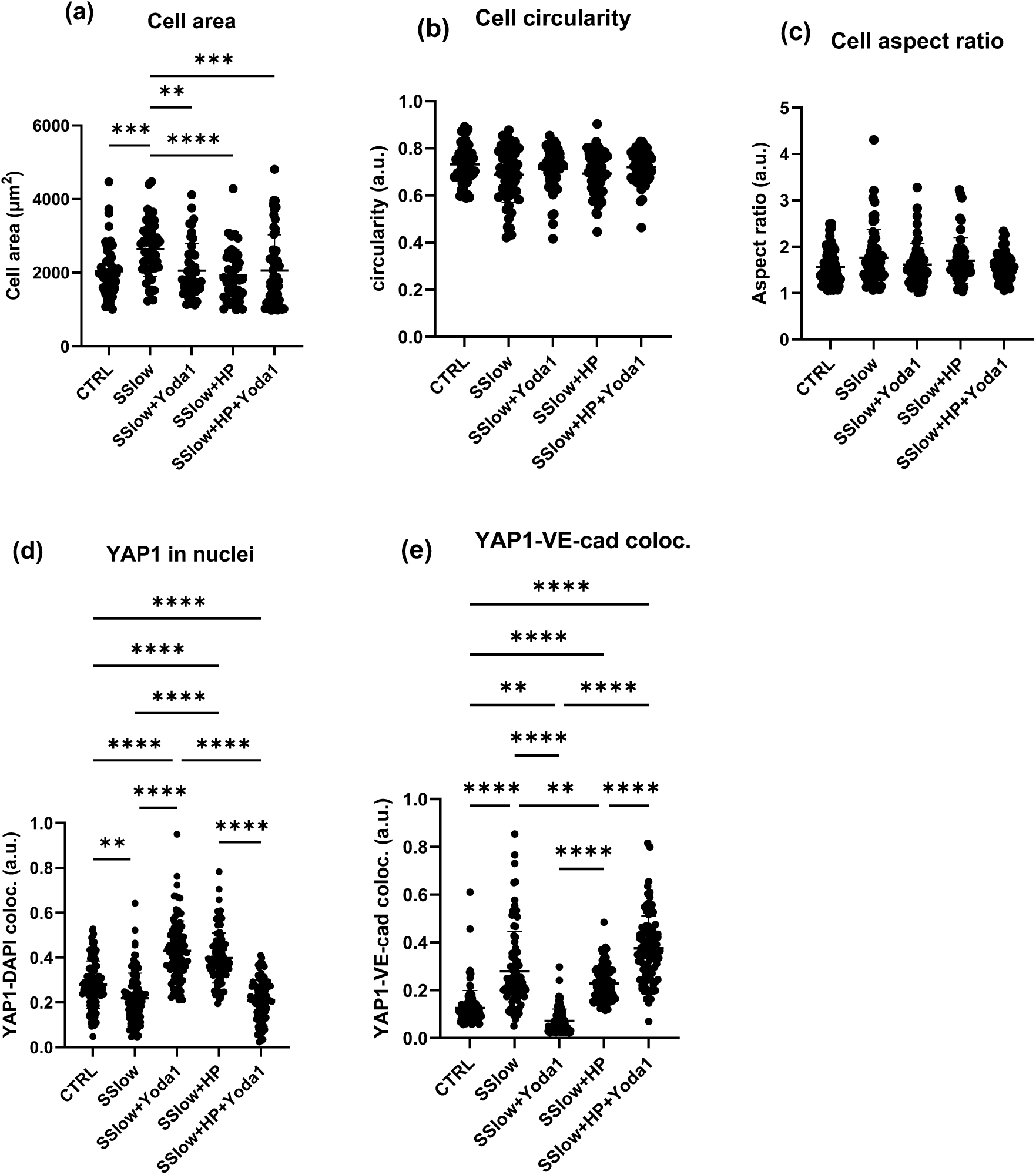
Exposure to elevated hydrostatic pressure increases YAP1 nuclear localisation, reduces association with VE-cadherin and alters VE-cadherin expression pattern at cell-cell contacts. HUVECs cultured within the microchannels were exposed to the different flow conditions with or without Yoda1 (2 µM) for 1 h after which the cells were fixed, permeabilised and immunostained for VE-cadherin (red), YAP1(green) and nuclei (blue). Representative immunofluorescence images of HUVECs treated to (A-D) SSlow only (E-H) SSlow only+Yoda1 (I-L) SSlow+HP (M-P) SSlow+HP+Yoda1 (Q-T) CTRL. (a) average cell area (b) cell circularity (c) cell aspect ratio (d) YAP1 nuclear localisation quantified by Manders’ Colocalization Coefficient (e) YAP1-VE-cadherin colocalisation quantified by Manders’ Colocalization Coefficient. Data represented as Mean±SD, N = 3, at least 50 cells were analysed per repeat. Flow direction: top to bottom. White arrows: linear VE-cadherin junctions, yellow arrows: disrupted linear VE-cadherin junctions, white arrow heads: mature VE-cadherin fingers, yellow arrow heads: immature VE-cadherin fingers. Scale = 20 µm. Mag = 40×

In comparison, SSlow+HP and SSlow+HP+Yoda1 demonstrated a serrated VE-cadherin pattern with prominent finger-like structures at the junctions as shown in Figure 1 (J) and (N) respectively. These VE-cadherin finger-like structures (referred to as VE-cadherin fingers henceforth) were more pronounced in the SSlow+HP+Yoda1 conditions (denoted by white arrow heads in Figure 1 (N)) compared to the immature VE-cadherin fingers seen in SSlow+HP conditions (denoted by yellow arrow heads in Figure 1 (J)). This result demonstrates the influence of Piezo-1 activation in mature VE-cadherin finger formation during elevated hydrostatic pressure application. In addition, these VE-cadherin fingers were not observed in the SSlow only+Yoda1 which shows that Piezo-1 activation alone is not sufficient for VE-cadherin finger formation and hence this effect signifies the distinct mechanobiological stimulus of elevated hydrostatic pressure. Although there was no difference in the cell circularity or aspect ratio between the different conditions as shown in Figure 1 (b) and (c) respectively, SSlow only treated cells demonstrated a larger cell area compared to the other conditions as shown in Figure 1 (a).

Compared to SSlow only, there was increased YAP1 nuclear colocalisation in both SSlow only+Yoda1 and SSlow+HP as shown in Figure 1 (d), suggesting an increase in YAP1 nuclear signalling in the latter conditions. Interestingly, SSlow only showed increased colocalisation of YAP1 and VE-cadherin compared to SSlow only+Yoda1 and SSlow+HP conditions as shown in Figure 1 (e). This suggests that in the SSlow only+Yoda1 and SSlow+HP conditions, there is an increased dissociation of YAP1 from VE-cadherin and hence increased accumulation of YAP1 in nuclei. Comparing the latter two conditions, the fact that SSlow+HP demonstrated higher YAP1-VE-cadherin colocalisation than SSlow only+Yoda1 demonstrates the distinct effect of hydrostatic pressure. It was also interesting to note that compared to SSlow only+Yoda1, the addition of Yoda1 to SSlow+HP conditions demonstrated a surprising increase in YAP1-VE-cadherin colocalisation and a corresponding reduction in the YAP1 nuclear colocalisation. This observation suggests that the mechanobiological stimuli of elevated hydrostatic pressure can significantly impact the Piezo-1 activation signalling effects compared to the shear stress alone conditions.

### 4.2 Elevated hydrostatic pressure leads to increased EPS8 cytoplasmic localisation

EPS8 is an adaptor protein that regulates cell-cell junctional dynamics by transiently binding to VE-cadherin during early junctional maturation, a stage characterized by high VE-cadherin turnover and inactive PI3K [51, 52]. In the current study, cells were exposed to the different flow conditions in the presence or absence Yoda1 (2 µM) and Wortmannin (1.25 µM), a PI3K enzyme inhibitor, for 1 h after which they were analysed for EPS8 cytoplasmic area as well as EPS8-VE-cadherin colocalisation.

When analysed for EPS8 cytoplasmic localisation, there was a significant increase in the EPS8 cytoplasmic area (average EPS8 positive immunostaining area outside the nuclei) in the SSlow+HP conditions compared to SSlow only and CTRL as shown in Figure 2 (a). Interestingly, SSlow+HP demonstrated significantly higher cytoplasmic EPS8-VE-cadherin colocalisation compared to both SSlow only and CTRL conditions as shown in Figure 2 (b).

**Figure 2.**
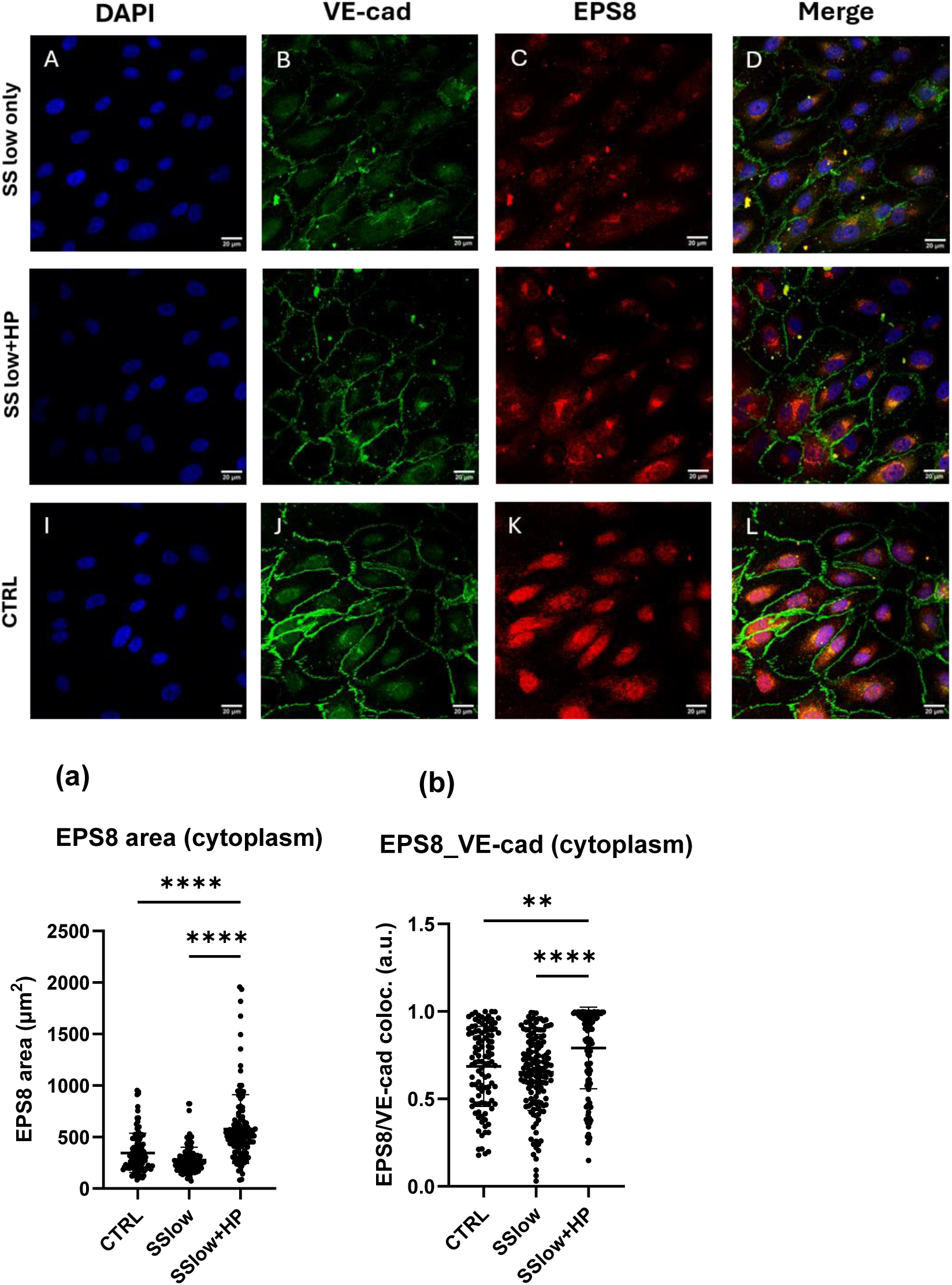
Exposure to elevated hydrostatic pressure led to increased cytoplasmic EPS8 area and promoted EPS8-VE-cadherin cytoplasmic colocalisation. HUVECs cultured within the microchannels were exposed to SSlow only and SSlow+HP for 1 h after which the cells were fixed, permeabilised and immunostained for VE-cadherin (green), EPS8 (red) and nuclei (blue). Representative immunofluorescence images of HUVECs treated to (A-D) SSlow only (E-H) SSlow+HP (I-L) CTRL. (a) average EPS8 cytoplasmic area (b) EPS8 and VE-cadherin colocalisation quantified by Manders’ Colocalization Coefficient. Data represented as Mean±SD, N = 3, at least 50 cells were analysed per repeat. Flow direction: top to bottom. Scale = 20 µm. Mag = 40×

### 4.3 Wortmannin promoted EPS8-VE-cadherin association which was reduced by Yoda1 addition at elevated hydrostatic pressure

Next, in order to investigate the role of Piezo-1 and PI3K enzyme on hydrostatic pressure mechanotransduction, the HUVECs were treated with Yoda1 (2 µM) and/or Wortmannin (1.25 µM) at elevated hydrostatic pressure. As shown in Figure 3 (a) and (b), the different combinations of Wortmannin and Yoda1 did not produce any significant difference in cell area or circularity. When analysed for VE-cadherin expression patterns, the addition of Wortmannin demonstrated a diffuse VE-cadherin pattern mainly concentrated in the cell cytoplasm and exhibiting a thin, disrupted membrane VE-cadherin pattern (denoted by white arrows in Figure 3 (B) and (F)) in both SSlow+HP+Wortmannin and SSlow only+Wortmannin conditions as shown in Figure 3 (B) and (F), respectively.

**Figure 3.**
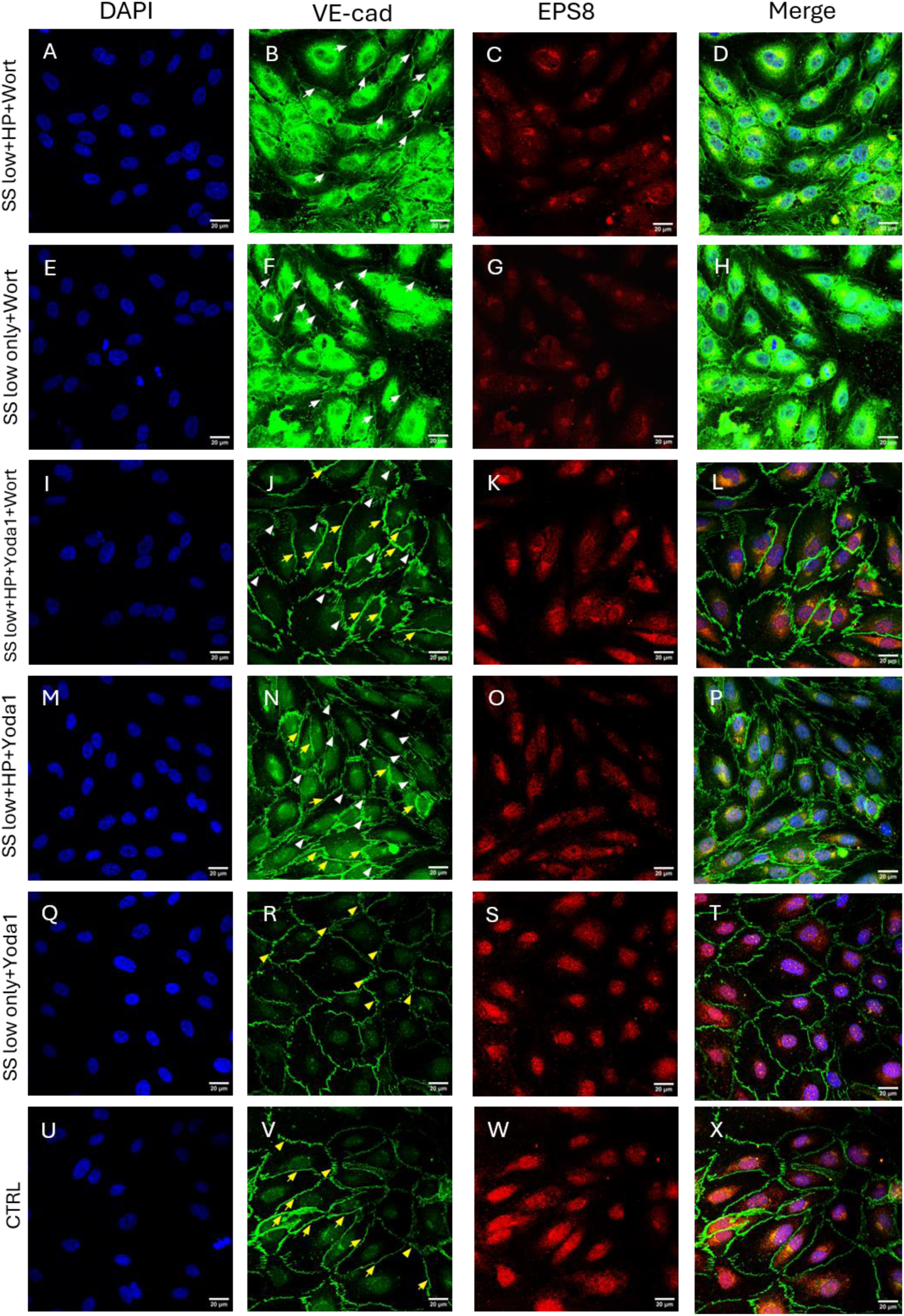
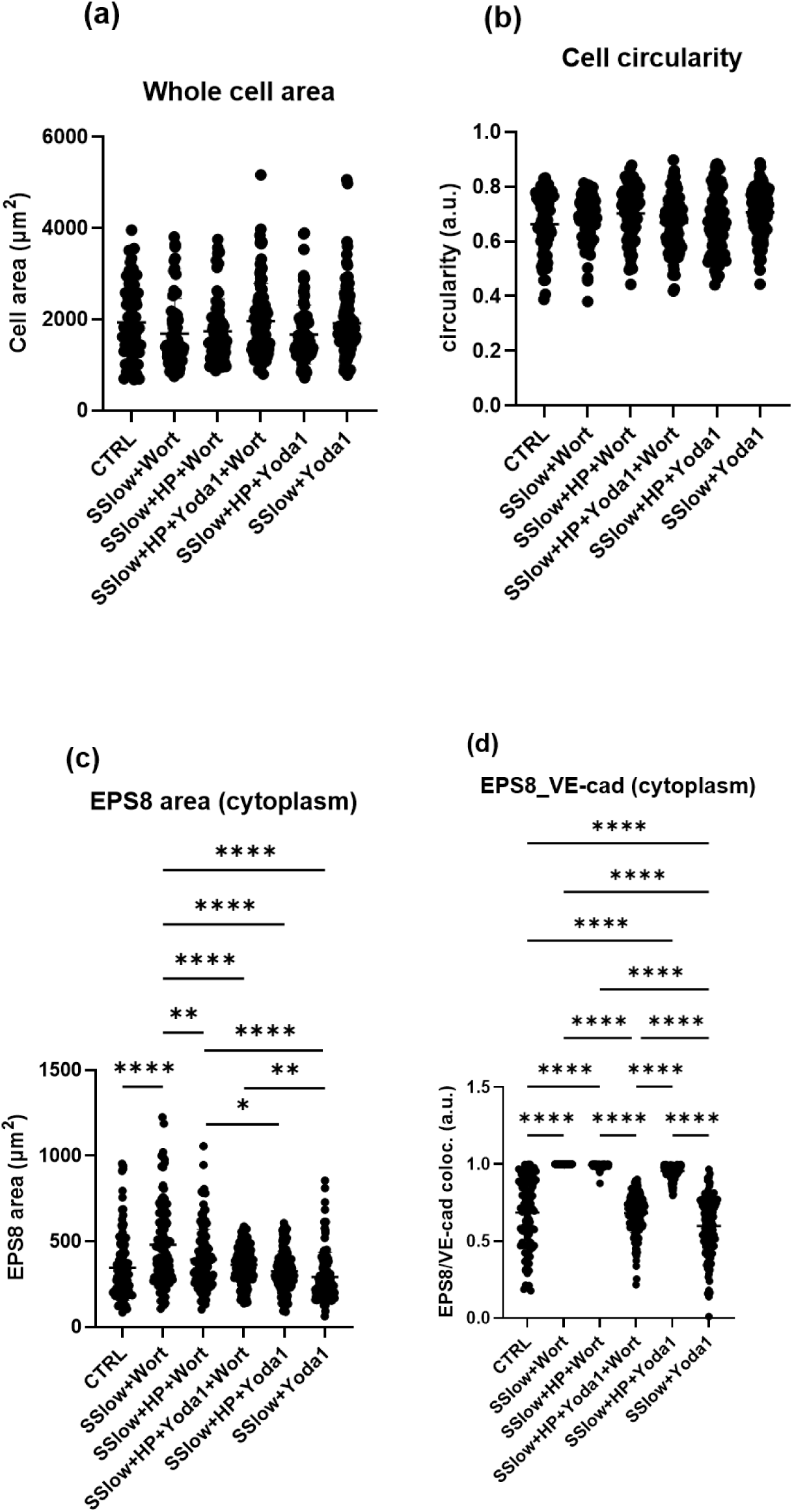
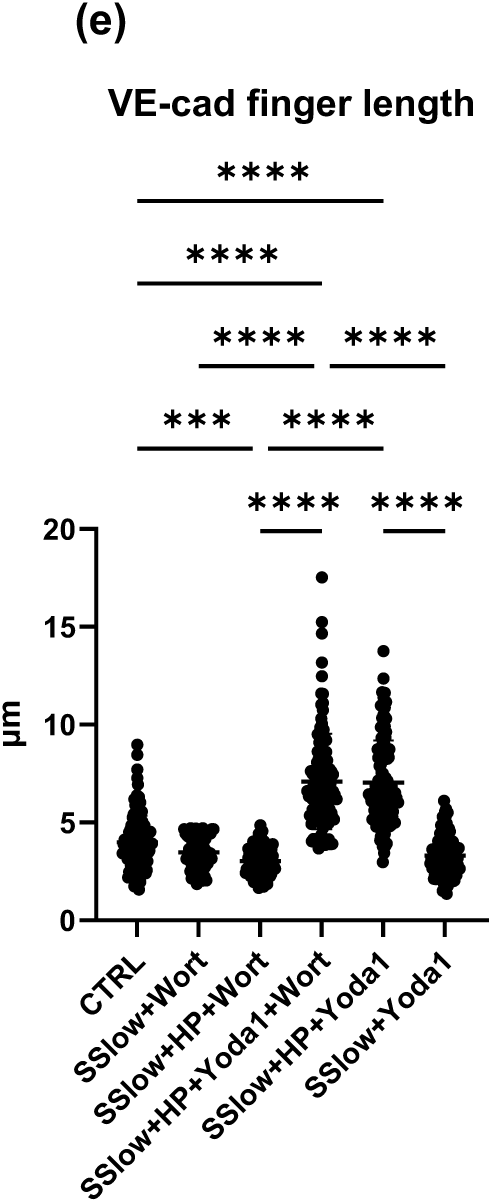
Wortmannin increased cytoplasmic EPS8 area and VE-cadherin colocalisation, decreased VE-cadherin finger length, while Yoda1 reversed these effects at elevated hydrostatic pressure. HUVEC cells were treated to the different flow conditions in the presence or absence of 1.25 μM Wortmannin (PI3K inhibitor) or 2 μM Yoda1 (piezo-1 agonist) or both for 1 h after which the cells were fixed, permeabilised and immunostained for VE-cadherin (green), EPS8 (red) and nuclei (blue). Representative immunofluorescence images of HUVECs treated to (A-D) SSlow+HP+Wortmannin (E-H) SSlow only+Wortmannin (I-L) SSlow+HP+Wortmannin+Yoda1 (M-P) SSlow+HP+Yoda1 (Q-T) SSlow only+Yoda1 (U-X) CTRL. (a) average cell area (b) cell circularity (c) average EPS8 cytoplasmic area (d) EPS8-VE-cadherin cytoplasmic colocalisation quantified by Manders’ Colocalization Coefficient (e) average VE-cadherin finger length. Data represented as Mean±SD, N = 3, at least 50 cells were analysed per repeat. Flow direction: top to bottom. White arrows: thin, linear VE-cadherin junctions, yellow arrows: thick, linear VE-cadherin junctions, white arrow heads: mature VE-cadherin fingers, yellow arrow heads: immature VE-cadherin fingers Scale = 20 µm. Mag = 40×

Despite there being no difference in the whole cell area between the different conditions as shown in Figure 3 (a), the EPS8 cytoplasmic area varied significantly between the different conditions as shown in Figure 3 (c). Interestingly, in both the Wortmannin only conditions (SSlow+Wortmannin, SSlow+HP+Wortmannin), there was a significant increase in both the cytoplasmic EPS8 area as well as EPS8-VE-cadherin cytoplasmic colocalisation compared to the conditions without Wortmannin as quantified in Figure 3 (c) and (d) respectively. Especially, SSlow+Wortmannin demonstrated increased cytoplasmic EPS8 area compared to all conditions. Interestingly, the addition of Yoda1 to SSlow+HP+Wortmannin exhibited a significant reduction in both the EPS8 cytoplasmic area and the EPS8-VE-cadherin cytoplasmic colocalisation as shown in Figure 3 (c) and (d) respectively, which demonstrates the potential role of Piezo-1 in regulating the EPS8 and VE-cadherin association in response to elevated hydrostatic pressure. Also interesting was the observation that SSlow+HP+Yoda1 demonstrated higher EPS8-VE-cadherin cytoplasmic colocalisation compared to SSlow only+Yoda1, thus demonstrating the distinct mechanobiological input of hydrostatic pressure in facilitating the downstream effects of Piezo-1 activation in regulating VE-cadherin junctions.

Further, Yoda1 addition to both the pressure conditions (SSlow+HP+Yoda1±Wortmannin) caused an increase in VE-cadherin finger length and exhibited thick, linear VE-cadherin junctions (denoted by yellow arrows in Figure 3 (J) and (N)) even in the presence of Wortmannin as shown in Figure 3 (e). These data indicate that despite PI3K inhibition, Yoda1 can preserve VE-cadherin assembly at cell-cell junctions in a PI3K-independent manner, albeit with prominent, mature VE-cadherin finger formation (denoted by white arrow heads in Figure 3 (J) and (N)) which have been reported to be a characteristic sign of endothelial junctional remodelling [67]. Also, similar to the YAP1 results, the influence of Piezo-1 activation on VE-cadherin finger formation was more pronounced in the elevated hydrostatic pressure conditions as SSlow only+Yoda1 treatment exhibited immature VE-cadherin fingers (denoted by yellow arrow heads in Figure 3 (R)) and were comparable to the Wortmannin and CTRL conditions. CTRL conditions predominantly exhibited thick, linear VE-cadherin conditions.

### 4.4 Elevated hydrostatic pressure caused increased endothelial permeability which was attenuated by Yoda1

In order to evaluate the endothelial monolayer barrier integrity in response to elevated hydrostatic pressure, HUVECs were cultured on biotinylated gelatine as mentioned in the Methods section 3.5 and were exposed to the different flow regimes after which Avidin-FITC (2.5 µg/mL) was introduced into the microfluidic channel. After 3 min incubation of Avidin-FITC at room temperature, the cells were washed to remove unbound Avidin-FITC after which they were fixed, stained for actin and imaged using confocal microscope. The mean F.I. of Avidin-FITC was calculated for each condition and normalised to the mean F.I. of the CTRL as described in the Methods section 3.5.

Comparing the different flow regimes, SSlow only and CTRL conditions demonstrated the lowest and comparable Avidin-FITC mean F.I as shown in Figure 4 (A) and (M) respectively and as quantified in Figure 4 (a). The addition of Yoda1 to SSlow only condition caused an ∼2.5-fold increase in Avidin-FITC permeability as shown in Figure 4 (D) and quantified in Figure 4 (a), thus demonstrating the permeability inducing effect of Piezo-1 activation in the shear stress alone conditions. A similar ∼2.45-fold increase in Avidin-FITC permeability was observed in the SSlow+HP condition as shown in Figure 4 (G) and quantified in Figure 4 (a) compared to the CTRL conditions. Interestingly, the addition of Yoda1 to the SSlow+HP condition caused a significant decrease in the Avidin-FITC permeability (∼1.57-fold change relative to the CTRL as quantified in Figure 4 (a)), despite exhibiting patches of high Avidin-FITC permeability (high permeability patches denoted by white arrows in Figure 4 (J)). This result demonstrates an interesting permeability inducing or protecting dual effect of Yoda1 depending on the presence or absence of elevated hydrostatic pressure mechanobiological stimuli.

**Figure 4.**
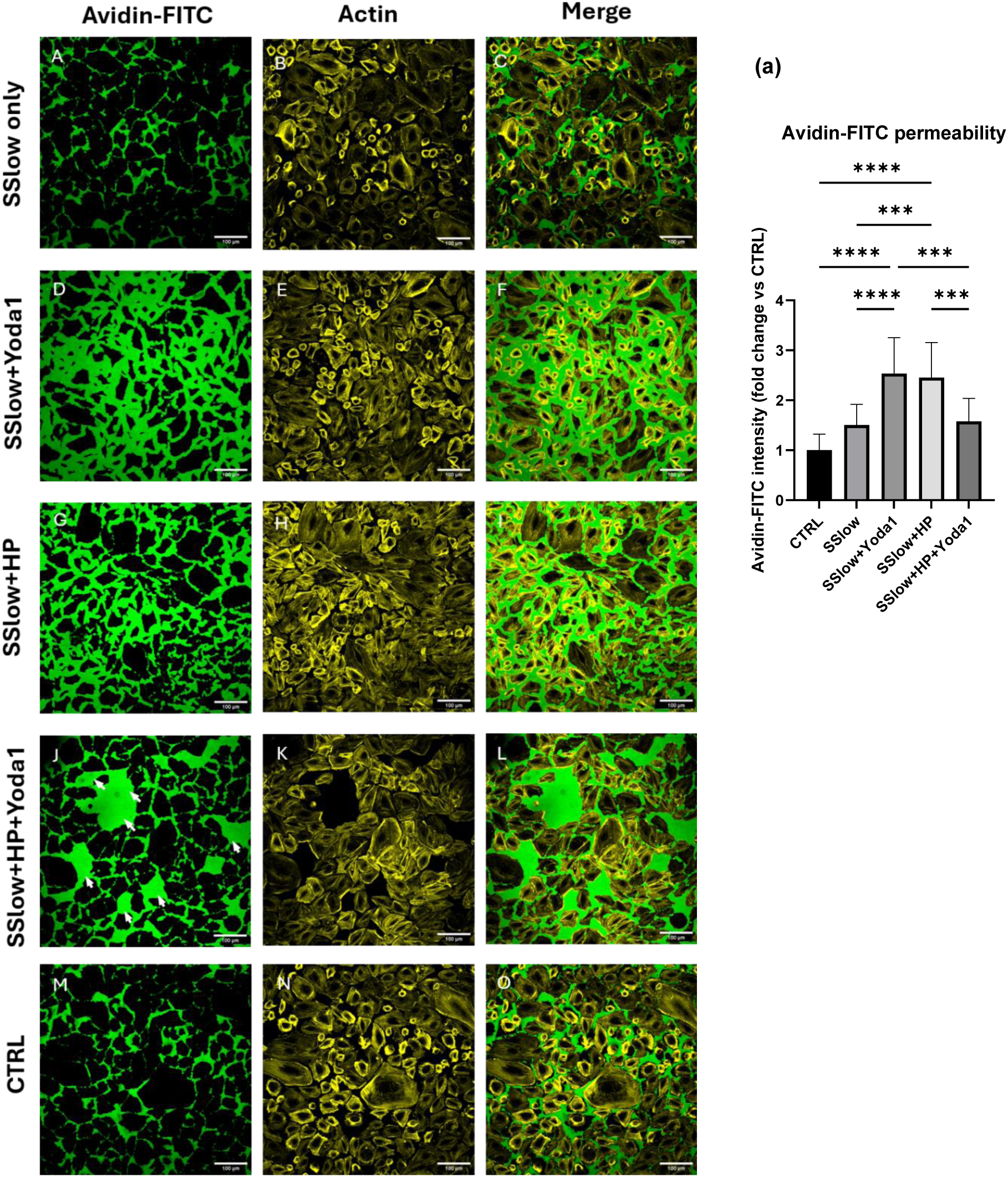
Yoda1 protects endothelial barrier integrity but only in the presence of elevated hydrostatic pressure and not in shear stress alone conditions. HUVECs cultured on biotinylated gelatine were treated to the different flow conditions for 1 h in the presence or absence of Piezo-1 agonist Yoda1 (2 µM) after which Avidin-FITC (2.5µg/mL) was added to the microchannel for 3 mins and incubated at room temperature. After washing the channels with DPBS to remove unbound Avidin-FITC, the cells were fixed and stained for actin using Phalloidin CF^®^594. The flow different conditions were (A-C) SSlow only (D-F) SSlow only+Yoda1(G-I) SSlow+HP (J-L) SSlow+HP+Yoda1 (M-O) CTRL. (a) Quantification of the Avidin-FITC permeability assay. Y-axis represented as normalised Avidin-FITC intensity. N = 3, five ROIs were analysed per sample per condition. Flow direction: top to bottom. White arrows: high permeability Avidin-FITC patches. Avidin-FITC (green), Actin (yellow). Scale = 100 µm.

### 4.5 Elevated hydrostatic pressure led to increased THP-1 adhesion which was abrogated by Yoda1

Endothelial barrier integrity and inflammatory activation are tightly coupled processes in the progression of microvascular dysfunction [68, 69]. To evaluate the functional impact of elevated hydrostatic pressure on endothelial inflammatory activation, we quantified THP-1 monocyte recruitment to HUVEC monolayers under the different flow regimes as detailed in Methods Section 3.6.

As shown in Figure 5 (A) and quantified in Figure 5 (a), SSlow only conditions demonstrated the least number of adhered monocytes (1.06-fold change vs CTRL) and were comparable to that of the CTRL conditions. The addition of Yoda1 to SSlow only resulted in a marginal increase in the number of adhered monocytes (1.65-fold change vs CTRL) although this increase was not statistically significant compared to SSlow only and CTRL. In comparison, exposure to elevated hydrostatic pressure resulted in a significant increase (∼3.6-fold change vs CTRL) in the number of adhered monocytes compared to both the shear stress only conditions (SSlow±Yoda1) and CTRL. Strikingly, the addition of Yoda1 to the SSlow+HP conditions attenuated this effect, resulting in a 2.2-fold reduction in adhered monocytes compared to the SSlow+HP conditions. This observed reduction in monocyte adhesion demonstrated the potential of Yoda1 to reduce recruitment of immune cells in elevated hydrostatic pressure conditions. As expected, treating the cells to a pro-inflammatory cytokine such as TNF-α elicited the most robust monocyte adhesion (∼5.1-fold vs CTRL) with an observed ∼1.5-fold increase compared to SSlow+HP.

**Figure 5.**
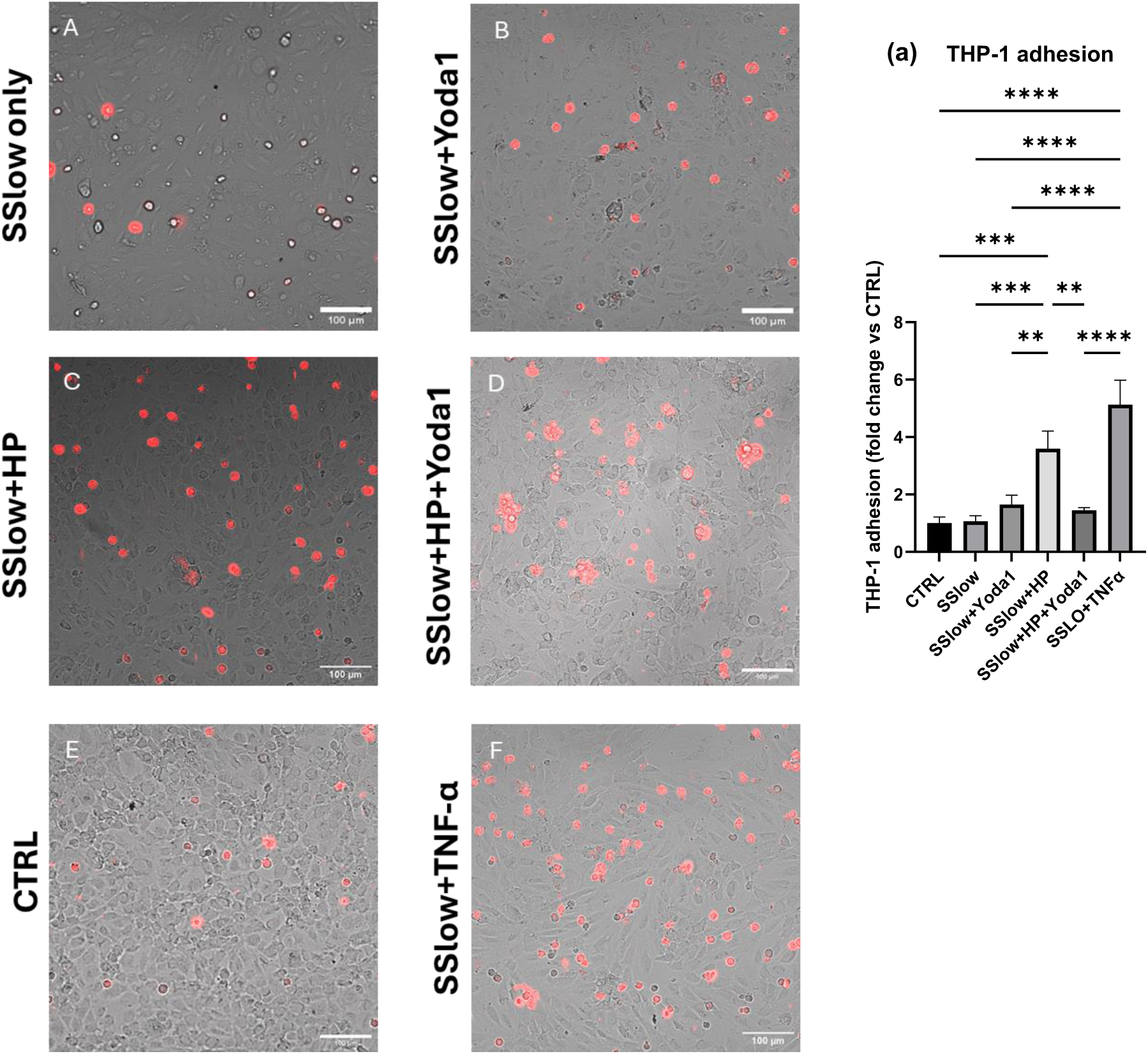
Elevated hydrostatic pressure promoted THP-1 adhesion which was attenuated by Yoda1 treatment. HUVECs cultured within the microfluidic device were treated to the different flow conditions with or without Yoda1 (2 µM) for 45 min after which THP-1 cells (200,000 cells/mL) were flowed over the endothelial monolayer for 15 min under the same flow conditions within a stage-top incubator. Time-lapse imaging was performed for the final 3 min of the experiment at a frequency of 0.5 Hz. The last frame of each time-lapse video was used to quantify the number of firmly adherent THP-1 cells. As a positive control, cells were treated to TNF-α (10 ng/mL) for 24 h prior to the perfusion of THP-1 at SSlow only flow conditions. The flow different conditions were (A) SSlow only (B) SSlow only+Yoda1 (C) SSlow+HP (D) SSlow+HP+Yoda1 (E) CTRL (F) SSlow+TNF-α. (a) Quantification of THP-1 adhesion to endothelial cells in the different flow conditions. Y-axis represented as normalised values relative to CTRL. N = 3, last frame of each experimental repeat was used to quantify THP-1 adhesion. Scale = 100 µm.

## 5 Discussion

Hydrostatic pressure is a fundamental mechanobiological force that regulates key cellular processes such as cell shape, volume, and proliferation in addition to controlling tissue level processes such as cell patterning, morphogenesis and growth [70, 71]. In vascular biology, although studies have predominantly focused on the influence of shear stress and cyclic mechanical stretch, the mechanisms of hydrostatic pressure mechanotransduction are poorly understood despite playing critical roles in regulating endothelial processes including actin remodelling and barrier integrity [72, 73], apoptosis [74], proliferation [75], endothelial tube formation [28] and angiogenesis [27]. Moreover, these studies investigating hydrostatic pressure mechanobiology have reported contradictory results [26, 76], possibly due to the varying experimental set-ups between the studies. Employing an easy-to-implement microfluidic platform, the current study investigated the combined impact of low shear stress and elevated hydrostatic pressure on endothelial junctional dynamics with a particular focus on VE-cadherin junctional expression pattern, its cytoplasmic colocalisation with YAP1 and EPS8, both of which are known to play a critical role in junctional stability. To our knowledge, this is the first study to investigate the roles of Piezo-1 activation and PI3K inhibition in the context of YAP1, VE-cadherin and EPS8 colocalisation at elevated hydrostatic pressure, thereby providing insight into a sparsely investigated elevated hydrostatic pressure-Piezo-1-PI3K mechanobiological axis.

When analysed for VE-cadherin junctional patterns, SSlow only+Yoda1 demonstrated intermittent disruption in linear VE-cadherin staining at cell-cell junctions while SSlow+HP and SSlow+HP+Yoda1 exhibited serrated, finger-like structures at the cell-cell contacts. Past studies have shown the impact of varying flow regimes [77] and prolonged exposure to shear stress [78] on VE-cadherin junctional expression wherein discontinuous or punctuated patterns were observed in response to mechanical or biochemical stimuli. Such discontinuous or punctuated patterns, although a transient phenomenon, have been reported to be indicative of remodelling junctions and are primarily mediated by actomyosin stress fibre formation and its association with junctional VE-cadherin [79, 80]. In our past study, we demonstrated that a short-term (1 h) exposure to elevated hydrostatic pressure at low shear stresses led to the formation of VE-cadherin fingers at cell-cell junctions similar to that observed in the SSlow+HP conditions of the current study [63]. Hayer et al (2016) demonstrated that such VE-cadherin fingers involve formation of engulfed plasma membrane tubes between the leader and follower cells during wound healing cell migration [67]. In another interesting study, Huveneers et al (2012) demonstrated that vinculin along with radial actin bundles are involved in the formation of remodelling VE-cadherin finger-like structures (which the study refers to as focal adhesion junctions) [81]. In this context, further investigation into the specific molecular processes driving the formation of these VE-cadherin fingers in response to elevated hydrostatic pressure may provide valuable insights into endothelial dysfunction mechanisms.

In order to perform vascular functions such as permeability, angiogenesis and inflammation, endothelial cell-cell junctions are dynamically reorganised in response to mechanical and biochemical stress [80]. In recent years, YAP1 has been identified as a key mechanotransducer in vascular pathogenesis [82]. In vivo studies have shown increased nuclear accumulation of YAP1 in the atheroprone regions of the mice arteries [83] and depletion of the *Yap* gene have been shown to cause decreased plaque formation in *ApoE^−/−^* mice [84]. Previous mechanistic studies have demonstrated that laminar flow promotes an atheroprotective endothelial phenotype by downregulating YAP1 nuclear accumulation [84, 85]. In the current study, while SSlow only demonstrated reduced nuclear accumulation of YAP1 comparable to that of the CTRL, application of hydrostatic pressure increased the nuclear accumulation of YAP1 and reduced YAP1-VE-cadherin colocalisation. Similarly, at elevated hydrostatic pressure, there was an increase in EPS8 cytoplasmic area and its cytoplasmic colocalisation with VE-cadherin, demonstrating enhanced association of EPS8 to VE-cadherin in the elevated pressure-induced adherens junction reorganisation. These results suggest that elevated pressure-induced changes in YAP1 localisation and enhanced EPS8-VE-cadherin colocalisation may contribute to junctional remodelling and downstream changes in endothelial permeability and monocyte adhesion. In one past study focusing on this mechanobiological axis, Giampietro et al (2015) reported that the association of EPS8 with VE-cadherin in remodelling junctions led to dephosphorylation of YAP1 and subsequent nuclear translocation whereas in stabilised junctions, phosphorylated YAP1 bound to VE-cadherin wherein EPS8 was excluded [52]. In this context, it would be interesting to study the phosphorylation dynamics of YAP1 in relation to EPS8-VE-cadherin colocalisation at elevated hydrostatic pressure in order to elucidate the specific mechanisms of junctional reorganisation and downstream effector functions. Following nuclear translocation, YAP1 binds to the TEAD transcription factors which activate junctional remodelling and pro-inflammatory genes such as ICAM-1, VCAM-1 promoting adhesion of leukocytes [86]. This correlates well with the current study wherein the application of hydrostatic pressure led to increased permeability of Avidin-FITC and increased adhesion of THP-1 monocytes compared to SSlow only conditions and CTRL, thus demonstrating the pro-permeability and pro-adhesive endothelial activation effects of elevated hydrostatic pressure. In agreement with our observation, one past study by Michell et al (2021) demonstrated that elevated intraluminal pressure led to increased adhesion of leukocytes with a corresponding increase in Intercellular adhesion molecule (ICAM-1) and monocyte chemoattractant protein 1 (MCP-1) [87].

In the current study, inhibiting PI3K using Wortmannin at elevated hydrostatic pressure demonstrated an increase in EPS8-VE-cadherin cytoplasmic colocalisation compared to the CTRL. One possible mechanism for this observed result could be that inhibiting PI3K promotes recruitment of EPS8 to the VE-cadherin junctions which renders the junction highly unstable [88]. Although a number of past studies have reported on the influence of the different PI3K isoforms in regulating VE-cadherin junctional clustering and barrier function [89, 90], only one past study has reported on the specific mechanobiological relationship between PI3K, EPS8 and VE-cadherin. In this study, the authors reported that the dissociation of EPS8 from VE-cadherin during endothelial remodelling phase resulted in increased activity of PI3K-Akt which prevented YAP1 nuclear translocation [52]. Another possible mechanism is that PI3K activity downregulates actomyosin contractility and hence when PI3K is inhibited, actomyosin tension increases, pulling the VE-cadherin adherens junctions apart [91]. Given the established role of EPS8 in both actin cytoskeletal remodelling [92] and adherens junction regulation [93, 94], it is plausible that EPS8 serves as an important intermediary linking mechanical stimuli to endothelial junctional dynamics. Therefore, it would be of considerable interest to investigate the mechanisms by which EPS8 regulates VE-cadherin and actin junctional remodelling and how these mechanisms are influenced by PI3K inhibition under elevated hydrostatic pressure conditions. Further, since our current study relied on pharmacological PI3K inhibition using Wortmannin, future work using isoform-specific inhibition or genetic perturbation will be required to define the precise PI3K isoforms involved in pressure-dependent VE-cadherin/EPS8 regulation.

Piezo-1 activation via Yoda1 in the SSlow only and SSlow+HP conditions exhibited an interesting opposing effect in YAP1 expression. While SSlow only+Yoda1 demonstrated increased YAP1 nuclear localisation and decreased YAP1-VE-cadherin colocalisation, similar to that of the SSlow+HP conditions, the SSlow+HP+Yoda1 conditions demonstrated a relative decreased YAP1 nuclear localisation and increased YAP1-VE-cadherin colocalisation. Surprisingly, the addition of Yoda1 in the SSlow+HP conditions also led to a marked reduction in both junctional permeability and THP-1 adhesion. These results indicate that Piezo-1 activation can potentially attenuate the elevated hydrostatic pressure induced barrier disruptive and pro-adhesive inflammatory effects.

Previous studies have shown Piezo-1 to be the key regulator in shear stress [38, 95, 96] and hydrostatic pressure mechanotransduction [63, 97, 98]. Taken together, these results suggest an interesting mechanism that Piezo-1 activation via Yoda1 under an already mechanically stimulated environment of elevated hydrostatic pressure conditions may not amplify the mechanical signal in a linear fashion but instead may exhibit an adaptive mechanism which shifts the cells towards a more stable and less monocyte-adhesive phenotype. One possible mechanism for this could be that sustained Piezo-1 mediated Ca²⁺ influx activates an enzyme called calpain which protects VE-cadherin at junctions by cleaving Src kinase, a molecule known to destabilise VE-cadherin at cell-cell junctions [99]. Stable VE-cadherin junctions increase YAP1 sequestration at junctions through VE-cadherin/α-catenin complexes and thereby suppressing YAP1 mediated pro-inflammatory endothelial activation [100, 101]. In line with this hypothesis, Wang et al (2025) reported on the inflammation suppressive and pro-angiogenesis effects of Piezo-1 activation via Ca^2+^/HIF-1α/VEGF signalling pathway towards ameliorating ischaemic reperfusion injury [36]. Other studies have shown Piezo-1 activation via Yoda1 to promote anti-inflammatory effects by increasing the expression of KLF2 or KLF4 [102] and decreasing NF-κB-dependent ICAM-1, VCAM-1 expression [103]. These studies may also explain another interesting observation of the current study that while SSlow+Yoda1 condition showed a significant increase in junctional permeability of avidin-FITC, there was no notable increase in THP-1 adhesion in the same condition compared to both SSlow only and CTRL conditions. This suggests that Yoda1 addition in shear stress alone conditions, in the absence of an additional mechanical stimuli such as elevated hydrostatic pressure, can lead to junctional remodelling but not necessarily promote a pro-adhesive inflammatory phenotype. These findings demonstrate that in addition to shear stress, the distinct mechanobiological stimuli of elevated hydrostatic pressure plays a key role in regulating the downstream effects of Piezo-1 mechanosensitive channel. Future studies employing Piezo-1-deficient endothelial models together with real-time calcium imaging and live-cell analysis of VE-cadherin dynamics under elevated hydrostatic pressure will be required to elucidate the precise molecular mechanisms governing elevated hydrostatic pressure-dependent endothelial junctional dynamics, barrier integrity and pro-adhesive inflammatory effects.

## 6 Conclusions

This study demonstrates that elevated hydrostatic pressure is a potent regulator of endothelial mechanobiology, promoting VE-cadherin finger-like junctional patterns, increased YAP1 nuclear localisation, altered EPS8-VE-cadherin cytoplasmic associations, enhanced endothelial permeability and THP-1 monocyte adhesion. PI3K inhibition in the elevated pressure conditions led to a decreased, thin VE-cadherin junctional pattern along with increased cytoplasmic EPS8-VE-cadherin colocalisation. Addition of Yoda1 to the PI3K inhibited conditions led to formation of thick VE-cadherin fingers at junctions along with reduced EPS8-VE-cadherin colocalisation, thus highlighting the co-ordinated action of Piezo-1 and PI3K signalling in elevated hydrostatic pressure-dependent junctional organisation. A key finding of this work was the context-dependent effect of Piezo-1 activation in endothelial cells. While Yoda1 treatment under shear stress alone demonstrated disrupted VE-cadherin junctional pattern, increased permeability, no impact on THP-1 adhesion, and increased YAP1 nuclear accumulation, Piezo-1 activation under combined shear stress and elevated hydrostatic pressure demonstrated VE-cadherin finger formation at junctions, decreased permeability and THP-1 adhesion, and decreased nuclear YAP1 localisation. These findings suggest that under an elevated hydrostatic pressure-loaded condition, Piezo-1 signalling may function as an adaptive regulator of endothelial barrier and pro-adhesive inflammatory responses. Collectively, this work identifies hydrostatic pressure as an important but underexplored regulator of endothelial dysfunction and identifies Piezo-1 and PI3K signalling as promising mechanobiological pathways involved in modulating VE-cadherin, YAP1, and EPS8 activity.

## 8 Data Availability

All data supporting the findings of this study are included in the manuscript and its Supplementary information document.

## 10 Author contributions

PVB designed, conducted experiments and analysed the data. TA performed the analytical solutions for flow parameters within the rectangular microfluidic channel used in this study. PVB wrote the manuscript, which was revised by DV, MJHS and LMG. All authors approved the final version of the manuscript.

## 11 Acknowledgements

This work was supported by funding received from the UK Engineering and Physical Sciences Research Council (EPSRC) Programme Grant PREMIERE (EP/T000414/1). The authors would like to thank Dr. Pradeep Kesavanarayana (University College London) and Dr. John James (University of Warwick) for their valuable critical discussions.

