## Supplementary Information for "Elevated hydrostatic pressure modulates endothelial junctional mechanotransduction through VE-cadherin remodelling and altered association with YAP1, EPS8: an endothelium-on-chip study"

### S1 Shear stress and pressure distributions within the channel

The wall shear stress (WSS) and the hydrostatic pressure distribution within the microfluidic channel were calculated using an analytical solution reported in our previous study (1). Syringe pumps (World Precision Instruments, Hitchin, UK) were used to achieve the desired shear stress for different conditions by adjusting the flow rate ($Q$). For a rectangular channel with width $2W$ and height $2H$, the velocity field can be written (2):

$$u(x,y) = -\frac{16c_1W^2}{\pi^3}\sum_{n=0}^{\infty}\frac{(-1)^n}{(2n+1)^3}\left[1-\frac{\cosh\left(\frac{(2n+1)\pi y}{2W}\right)}{\cosh\left(\frac{(2n+1)\pi H}{2W}\right)}\right]\cos\left(\frac{(2n+1)\pi x}{2W}\right) \quad (1)$$

Where $c_1$ depends on the average velocity, $u_m = Q/(2W \times 2H)$, and can be written as:

$$c_1 = -3\frac{u_m}{W^2}\frac{1}{1-\frac{192}{\pi^5}\frac{W}{H}\sum_{n=0}^{\infty}\frac{1}{(2n+1)^5}\tanh\left(\frac{(2n+1)\pi H}{2W}\right)} \quad (2)$$

From this, WSS at the bottom wall of the channel (where the cells are cultured) is written as:

$$\mu\frac{\partial u(x,y)}{\partial y}\Big|_{y=-H} = -\mu\frac{16c_1W^2}{\pi^3}\sum_{n=0}^{\infty}\frac{(-1)^n}{(2n+1)^3}\left[\frac{(2n+1)}{2W}\frac{\sinh\left(\frac{(2n+1)\pi H}{2W}\right)}{\cosh\left(\frac{(2n+1)\pi H}{2W}\right)}\right]\cos\left(\frac{(2n+1)\pi x}{2W}\right) \quad (3)$$

Elevated hydrostatic pressure (HP) conditions were achieved by raising the outlet tube to 40 cm above the inlet as shown in Figure S1 (A, B). At this height, the pressure drop at the channel outlet ($P_{out}$) can be derived from:

$$P_{out} = P_{atm} + \rho g h + \frac{\mu Q L}{32\pi D^4} \quad (4)$$

where $P_{atm}$ is the atmospheric pressure, $\rho$ and $\mu$ are the density (1,000 kg $m^{-3}$) and dynamic viscosity (0.00072 Pa s at 37°C) of the growth medium, $h$ is the height of the outlet tubing (40 cm). Pressure drop due to friction is represented by $\frac{\mu QL}{32\pi D^4}$ wherein $Q$ is the flow rate, $L$ is the length of the channel and $D$ is the inner diameter (1 mm) of the outlet tubing. Given the very low flow rates, the friction pressure drop is negligible (<1 Pa) when compared to the atmospheric pressure. Hence the pressure drop at the outlet is $P_{out} - P_{atm} = 3924\ Pa$ (29.43 mmHg). In vascular conditions, high compartmental pressure (≥30 mmHg) can compress the venous blood vessels first (3-5), owing to their low-pressure values (4-10 mmHg). hence we wanted to investigate the impact of this elevated pressure (30 mmHg) on the endothelial cells of the venules.

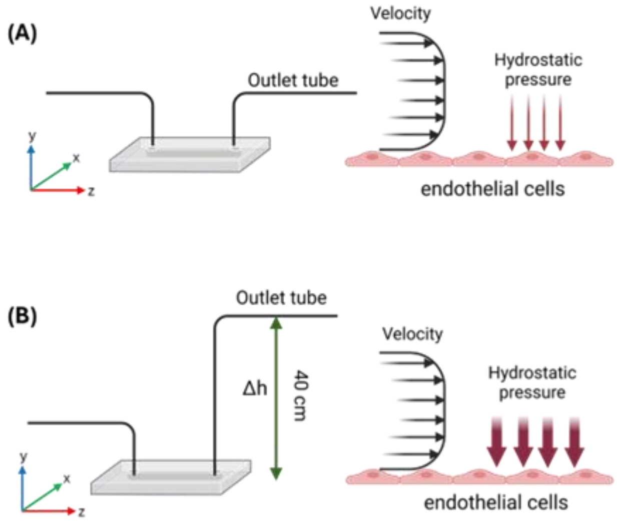


**Figure S1. (A)** Schematic of the microfluidic channel perfusion during normal flow conditions and **(B)** during application of elevated hydrostatic pressure conditions wherein the height of the outlet tube was elevated by 40 cm relative to the inlet tube, thereby maintaining the same shear stress profile but increasing the pressure experienced by the cells (x,y,z = W, H, L).

For a rectangular channel, the pressure drop along the channel ($z$-axis) can be expressed (2, 6):

$$\frac{dp}{dz} = \frac{\lambda \rho u_m^2}{2D_h} \qquad (5)$$

where $D_h$ is the hydraulic diameter and $\lambda$ depends on the aspect ratio of the channel ($\alpha = W/H$) and the Reynolds number ($Re = \rho u_m D_h/\mu$):

$$\lambda = \frac{96}{R_e}(1 - 1.3553\alpha^{-1} + 1.9467\alpha^{-2} - 1.7012\alpha^{-3} + 0.9564\alpha^{-4} - 0.2537\alpha^{-5}) \quad (6)$$

The pressure at a given point in the channel, with $P_{out}$ being the pressure at the exit of the channel, $L$ being the channel length and $z$ being the position from the inlet (i.e. $L{-}z$ is the position in the channel relative to the outlet) is given by:

$$p(z) = P_{out} + (L - z) * \frac{dp}{dz} \qquad (7)$$

For SSlow only conditions, the $P_{out} - P_{atm}$ at inlet $p$ ($z$ = 0) is 47.1 $Pa$ and the $P_{out} - P_{atm}$ at the outlet $p$ ($z$ = 17) is 0. Similarly, for the SSlow+HP conditions, $P_{out} - P_{atm}$ at the inlet

$p$ ($z$ = 0) for SS low+HP 3971.18 $Pa$ respectively and at the outlet $p$ ($z$ = 17), $P_{out} - P_{atm}$ is 3924 $Pa$. Hence the pressure difference (ΔP) along the channel is 47.1 $Pa$ respectively for both conditions. The full range of the $P_{out} - P_{atm}$ values at every point of $p(z)$ along the channel for both the SSlow as well as SSlow+HP conditions can be accessed via GitHub (https://github.com/thomas-abadie/rectangular-microchannel_velocity-pressure).
